# Integrated Molecular and AI-Based Diagnostics for Banana Diseases: Development, Optimization, and Field Deployment of LAMP and Computer Vision Technologies

**DOI:** 10.64898/2026.02.17.706421

**Authors:** Rimnoma S. Ouedraogo, Peter McCloskey, Beatrice Mwaipopo, Bipana Paudel Timilsena, Fatma H. Kiruwa, Milsort J. Kemboi, Mpoki M. Shimwela, David P. Hughes

## Abstract

Banana and plantain (*Musa* spp.) production in Sub-Saharan Africa is severely constrained by multiple diseases, with Banana bunchy top virus (BBTV) representing the most devastating viral threat. Inadequate diagnostic infrastructure limits effective management, particularly for asymptomatic infections disseminated through informal planting material exchange. This study presents an integrated diagnostic framework combining Loop-Mediated Isothermal Amplification (LAMP) molecular diagnostics with deep learning-based computer vision for rapid, scalable disease detection under field conditions. A LAMP assay targeting the BBTV DNA-S coat protein gene was developed using conserved sequences from diverse African isolates and validated with a simplified alkaline extraction protocol eliminating conventional DNA purification. The assay achieved 100% specificity and concordant detection with PCR and qPCR, reducing diagnostic time from 4 to 6 hours to 60 minutes. In-house recombinant *Bst* LF polymerase production demonstrated comparable enzymatic performance to commercial alternatives, with projected per-reaction cost reductions of 70 to 80%. Concurrently, an SSDLite MobileNetV2 object detection model was developed through 19 iterative training cycles on 19,914 field-collected images spanning 22 disease and physiological stress classes. The final model achieved recall rates of 92.5% for BBTV, 91.0% for Banana Xanthomonas Wilt, and 98.1% for healthy leaf classification, deployed via the PlantVillage mobile application for real-time offline diagnostics. A QR code-based metadata system integrates phenotypic AI assessments with molecular confirmation for comprehensive surveillance. This complementary framework addresses broad-scale phenotypic screening and molecular confirmation of pre-symptomatic infections, providing accessible tools to safeguard food security across Sub-Saharan Africa.

## 1.0 Introduction

Bananas and plantains (*Musa* spp.) are vital staple crops in Sub-Saharan Africa, providing year-round food sources that support food security and rural livelihoods across the continent. The region produces 30–33 million metric tons annually, sustaining millions who depend on these crops for daily nutrition (Scott, 2021; van Ittersum et al., 2016). However, yields remain far below their potential, averaging 5–30 metric tons per hectare compared to achievable yields exceeding 60 metric tons under optimal conditions (Smithson et al., 2001; Wairegi et al., 2010), a gap attributed to biotic and abiotic stresses, inadequate agronomic practices, and limited access to improved planting materials (Scott, 2021).

Among biotic constraints, Banana Bunchy Top Disease (BBTD), caused by Banana Bunchy Top Virus (BBTV), is recognized as the most devastating viral threat to banana production globally (Bouwmeester et al., 2023). First identified in Fiji in 1880 and reported in Africa in 1901, BBTD has been documented in more than 20 African countries (Dale, 1987; Kumar et al., 2011; Magee, 1927). BBTV is disseminated through two primary pathways: the movement of infected planting material, which facilitates long-distance spread, and local vector transmission by *Pentalonia nigronervosa*, the principal aphid vector (Watanabe et al., 2013).

Other significant diseases further threaten banana production across the region. Banana Xanthomonas Wilt (BXW), caused by *Xanthomonas vasicola* pv. *musacearum*, ranks among the most devastating bacterial diseases (Blomme et al., 2017; Ocimati et al., 2022). Fusarium wilt Tropical Race 4 (TR4) poses an existential threat following uncontained spread in Mozambique (Ploetz, 2015; Westerhoven et al., 2022). Sigatoka diseases, comprising Black Sigatoka (*Pseudocercospora fijiensis*) and Yellow Sigatoka (*Pseudocercospora musicola*), significantly impair photosynthetic capacity (Churchill, 2011; Kimunye et al., 2019). Arthropod pests compound these pressures: the banana weevil (*Cosmopolites sordidus*) causes 25–75% annual yield losses (Gold et al., 2001), while plant-parasitic nematodes, particularly *Radopholus similis*, inflict substantial root and corm damage (Speijer & Kajumba, 2000). Together, these biotic stresses create complex interactions that severely limit productivity (Blomme et al., 2013; Geberewold, 2019).

Despite these threats, diagnostic capacity in Sub-Saharan Africa remains inadequate. Conventional BBTV detection methods, including ELISA and PCR, require laboratory infrastructure and trained personnel unavailable in rural settings, a gap particularly consequential given BBTV’s ability to remain latent in planting material and spread through informal propagule exchange (Boonham et al., 2014; Kumar et al., 2015).

This study develops an integrated diagnostic framework combining artificial intelligence (AI)-based computer vision with molecular diagnostics. An AI model deployed on smartphones detects BBTD, BXW, and Sigatoka diseases from field images, enabling rapid phenotypic screening without specialized equipment. In parallel, a Loop-Mediated Isothermal Amplification (LAMP) assay provides cost-effective, field-deployable molecular detection of BBTV in asymptomatic planting material. These complementary technologies address distinct diagnostic needs: broad-scale phenotypic surveillance and molecular confirmation of pre-symptomatic infections, together providing scalable, accessible solutions for banana disease management and food security across Sub-Saharan Africa.

## 2.0 Materials and Methods

### 2.1 Molecular Diagnostics for BBTV Detection

#### 2.1.1 Sample Collection and Preparation

The samples were collected to comprehensively validate the specificity of the LAMP assay across diverse sample types. The validation panel comprised three categories: (i) BBTV-positive samples, including leaf and pseudostem tissues collected from symptomatic plants in the Kigoma and Morogoro regions of Tanzania, areas with documented active outbreaks (Mahuku & Kumar, 2023; Shimwela et al., 2022); (ii) negative controls from certified virus-free plants maintained under quarantine conditions; and (iii) field samples of unknown infection status, including both symptomatic and asymptomatic plants, to evaluate detection of different stage of infections critical for preventing BBTV spread through planting material (Németh & Kovács, 2025).

All sample collection followed established protocols for plant virus diagnostics to minimize cross-contamination (Boonham et al., 2008).). Leaf tissue was collected by punching a single disc from the third fully expanded leaf from the plant apex, excluding the youngest undeveloped “cigar leaf” (Nelson, 2006; Thomas, 2008). Sampling was conducted using sterile instruments, and collected tissues were immediately placed in individually labeled tubes and transported on ice to preserve integrity. To enable direct performance comparison across molecular diagnostic platforms, identical sample sets were evaluated using the LAMP assay, conventional PCR, and qPCR.

#### 2.1.2 Loop-Mediated Isothermal Amplification (LAMP) Assay Development

##### LAMP Primer Design and Validation

LAMP primers targeting the DNA-S component (coat protein gene) of BBTV were designed using the NEB LAMP Primer Design Tool Version 1.4.2 (https://lamp.neb.com/#!/). The primer design used a comprehensive dataset of 33 BBTV DNA-S sequences from diverse isolates, obtained from GenBank, ensuring the broad geographical representation and genetic diversity required for robust diagnostic assay development (Kumar et al., 2011).

Four specific primers were designed following the standard LAMP methodology: two outer primers (F3 and B3) and two inner primers, forward inner primer (FIP) and backward inner primer (BIP) (Table 1), which recognize six distinct regions of the target DNA sequence (Németh & Kovács, 2025; Notomi et al., 2000). This four-primer LAMP system provides the high specificity required for reliable BBTV detection while maintaining the simplicity needed for field deployment (Kim et al., 2019).

**Table 1.**
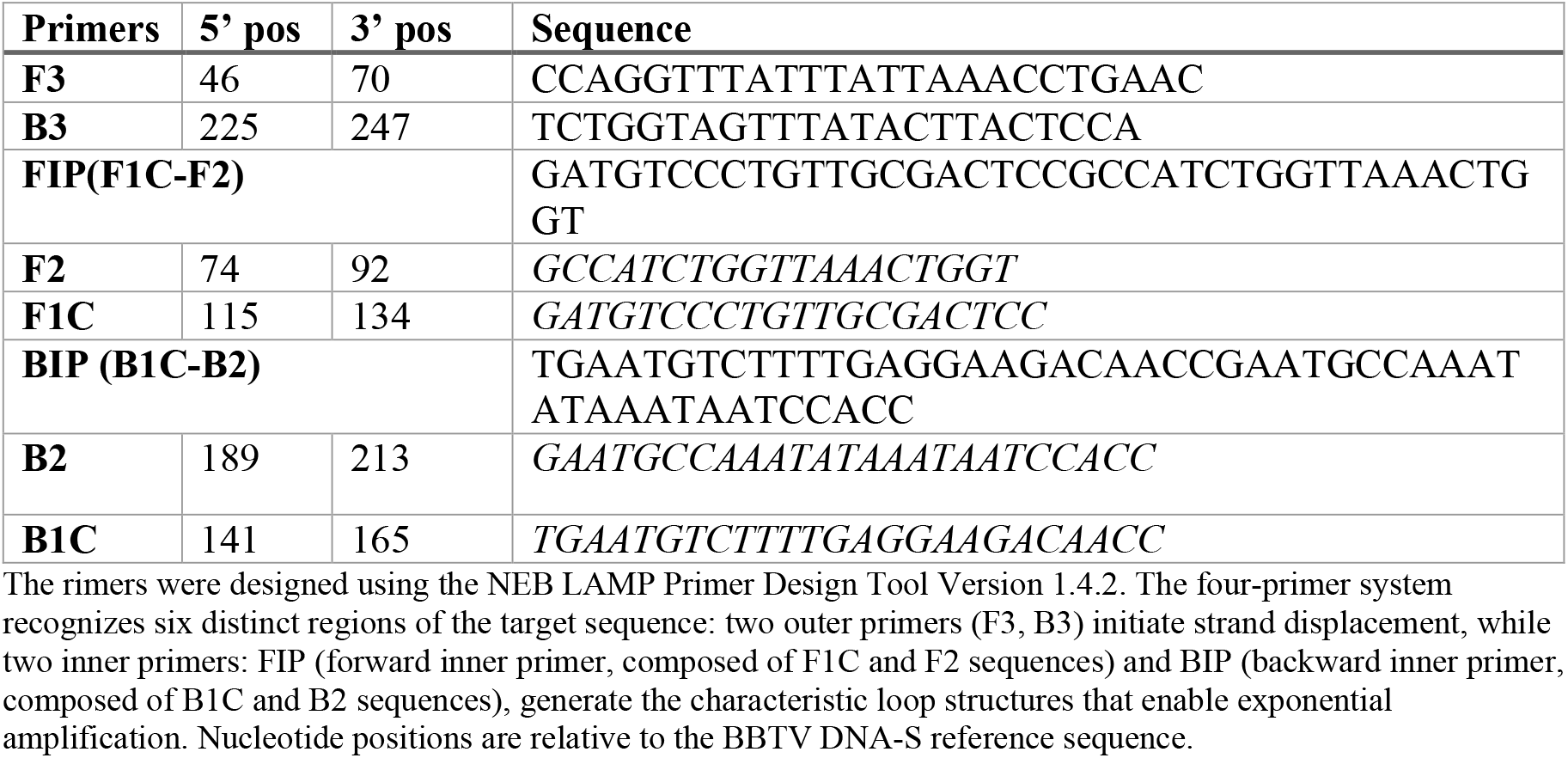
LAMP primer sequences targeting the DNA-S component (coat protein gene) of BBTV.

##### Recombinant Bst DNA Polymerase Expression and Purification

Recombinant Bst LF DNA polymerase was expressed using the pET21a(+)-Bst LF-6xHis expression vector (Addgene plasmid #159148), a gift from Andrea Pauli (RRID: Addgene_159148). Protein expression was performed in E. coli BL21(DE3)-Rosetta™ 2 competent cells (Novagen) (de Souza et al., 2023).

The purification protocol was adapted from established methods with modifications to optimize enzyme activity and stability (de Souza et al., 2023). Briefly, cells were grown in LB medium containing ampicillin (100 μg/mL) and chloramphenicol (34 μg/mL) at 37°C until the OD_600_ reached 0.6, then induced with 0.5 mM IPTG at 37°C for 4 hours. Cell lysis was performed using sonication in binding buffer (20 mM Tris-HCl, pH 8.0, 500 mM NaCl, 20 mM imidazole), and the enzyme was purified using Ni-NTA affinity chromatography. The purified enzyme was dialyzed against storage buffer (50 mM Tris-HCl, pH 8.0, 100 mM NaCl, 1 mM DTT, 50% glycerol). The activity was evaluated using NEB Bst-LF as a reference and stored at 8,000 U/ml.

##### Sample Preparation for LAMP Assay

A simplified sample-preparation method was developed to enable field deployment without requiring DNA extraction kits, following established protocols for the direct detection of plant viruses (Kim et al., 2019; Németh & Kovács, 2025). The method was adapted from (Chomczynski & Rymaszewski, 2006), with modifications to optimize DNA release from banana leaf tissue while maintaining compatibility with LAMP amplification.

A small disc (∼5 mm diameter) of banana leaf tissue was placed into a 1.5 mL microcentrifuge tube containing 980 µL of alkaline extraction buffer (2 mM NaOH; 6% polyethylene glycol-200). The leaf tissue was homogenized using a sterile toothpick to ensure thorough mixing with the buffer and complete cell disruption. After homogenization, 20 µL of neutralization buffer (0.1 M Tris-HCl, pH 8.0) was added to the tube and mixed by inversion. The mixture was allowed to sit for 2 minutes before proceeding to the LAMP assay to ensure DNA stabilization and pH neutralization (Chomczynski & Rymaszewski, 2006).

##### LAMP Reaction and Conditions

LAMP reactions were performed according to established protocols for plant virus detection, using a four-primer system (Notomi et al., 2000; Yeni et al., 2024). Each 12.5 μL reaction contained: 8U of Bst LF DNA polymerase, 1X Isothermal Amplification Buffer (20 mM Tris-HCl pH 8.8, 10 mM (NH_4_)_2_SO_4_, 10 mM KCl, 2 mM MgSO_4_, 0.1% Triton X-100), 6 mM additional MgSO_4_, 1.4 mM dNTPs (dATP, dGTP, dCTP, dTTP), 1.6 µM each of inner primers (FIP/BIP), 0.2 µM each of outer primers (F3/B3), 1X NEB LAMP Fluorescent Dye for real-time detection, and 2 µL of prepared sample supernatant. Nuclease-free water was added to achieve the final reaction volume (Kim et al., 2019; Németh & Kovács, 2025).

Negative controls consisted of 2 µL of extraction buffer alone or 30 ng of DNA extracted from certified healthy banana leaf samples. Positive controls contained known BBTV-positive samples validated by conventional PCR. The LAMP assay was performed on a Quantabio qPCR system (Quantabio, Beverly, MA, USA) under isothermal conditions at 65°C for 60 minutes, with fluorescence readings recorded every minute to monitor amplification in real time. A final inactivation step was performed at 85°C for 2 minutes to terminate the reaction and denature the polymerase (Notomi et al., 2000).

#### 2.1.3 Comparative Molecular Assays

##### DNA Extraction for PCR and qPCR

Total DNA of each sample was extracted using a modified cetyltrimethylammonium bromide (CTAB) protocol optimized for plant tissues containing high levels of polyphenolic compounds and polysaccharides (Schenk et al., 2023). The extraction buffer was supplemented with 2% polyvinylpyrrolidone (PVP) to remove phenolic compounds and 2.5% β-mercaptoethanol to prevent oxidation.

Briefly, 100 mg of leaf tissue was ground and homogenized in 750 μL of modified CTAB buffer (2% CTAB, 1.4 M NaCl, 0.2% β-mercaptoethanol, 2% PVP, 20 mM EDTA, 100 mM Tris-HCl, pH 8.0). The mixture was incubated at 65°C for 30 minutes with periodic vortexing. DNA was purified using chloroform: isoamyl alcohol (24:1) extraction followed by isopropanol precipitation. DNA pellets were washed with 70% ethanol, air-dried, and resuspended in TE buffer. DNA concentration was quantified using a NanoDrop spectrophotometer, and 30 ng of total DNA was used for each PCR and qPCR reaction.

##### Conventional PCR Amplification

Conventional PCR was performed using established BBTV-specific primers targeting the DNA-R component: BBT1 forward (5’-CTCGTCATGTGCAAGGTTATGTCG-3’) and BBT2 reverse (5’-GAAGTTCTCCAGCTATTCATCGCC-3’), designed to amplify a 349 bp fragment as previously described (Lestari & Hidayat, 2020). Reactions were prepared using OneTaq Quick-Load Master Mix (New England Biolabs, Ipswich, MA, USA) according to the manufacturer’s procedure.

Each 25 μL reaction contained: 12.5 μL OneTaq Quick-Load Master Mix, 0.5 μM each of forward and reverse primers, 24 ng template DNA, and nuclease-free water to final volume. Thermal cycling conditions were optimized as follows: initial denaturation at 94°C for 5 minutes, followed by 35 cycles of denaturation at 94°C for 30 seconds, annealing at 55°C for 45 seconds, and extension at 68°C for 1 minute, with final extension at 68°C for 6 minutes. PCR products were analyzed by electrophoresis on 1% agarose gels stained with ethidium bromide and visualized under UV illumination using a transilluminator (Lestari & Hidayat, 2020).

##### Quantitative Real-Time PCR (qPCR)

qPCR assays were performed using BBTV-specific primers targeting the DNA-R region: forward primer (5’-AAGGTCCCTTCGAGTTTGGT-3’) and reverse primer (5’-CAGCTATTCATCGCCTTCGT-3’) as described by Mendoza et al. (2024). Reactions were prepared using Luna Universal qPCR Master Mix (New England Biolabs, Ipswich, MA, USA) and performed on a QuantaBio qPCR system (95900-2C Q-qPCR instrument, Quantabio, Beverly, MA, USA).

Each reaction contained: 10 μL Luna Universal qPCR Master Mix, 0.25 µM each of forward and reverse primers, 25 ng template DNA extracted from positive samples and negative controls, and sterile deionized water to a final volume of 20 μL. Thermal cycling conditions were optimized as follows: initial denaturation at 95°C for 2 minutes (hold), followed by 40 cycles of denaturation at 95°C for 15 seconds, annealing at 60°C for 15 seconds, and extension at 72°C for 1 minute. Fluorescence readings were collected after each extension step to monitor amplification in real time (Mendoza et al., 2024).

### 2.2 Computer Vision-Based Disease Detection

#### 2.2.1 Dataset Collection and Curation

A comprehensive dataset was assembled with multiple banana diseases prevalent in Sub-Saharan Africa, including Banana Bunchy Top Disease (BBTD), Black Sigatoka (*Pseudocercospora fijiensis*), Yellow Sigatoka (*Mycosphaerella musicola*), Banana Xanthomonas Wilt (BXW) (*Xanthomonas vasicola* pv. *musacearum*),and physiological stress conditions such as nutrient deficiencies and tattered leaves. Healthy plant samples showing the purple color observed during the growth stages were also included to contrast with diseased and stressed crops and to enable the model to distinguish healthy crops (Ramcharan et al., 2019; Selvaraj et al., 2019).

Images were captured using Android smartphones equipped with cameras of a minimum 16-megapixel resolution to ensure sufficient image quality for feature extraction and disease symptom recognition (Mohanty et al., 2016a). All images were saved in JPEG or PNG formats with metadata preserved. Special emphasis was placed on capturing all stages of disease symptoms to ensure consistent recognition accuracy (Ramcharan et al., 2019).

#### 2.2.2 Data Processing and Quality Control

The collected images underwent systematic quality-control preprocessing, during which unsuitable images were identified and removed, including duplicates, blurred images, images with inadequate or excessive lighting, and images with occluded subjects (Barbedo, 2018; Hughes & Salathe, 2016). To ensure the model’s effectiveness under real-world field conditions, the training dataset included banana plants infected with single diseases as well as plants exhibiting co-infections with multiple pathogens. The curated images were then organized using a three-level hierarchical structure: first by disease category, second by plant anatomical focal point (leaf, stem, fruit, whole plant, buds), and third by country of collection, ensuring balanced representation across disease types, symptomatic locations, and geographic regions.

#### 2.2.3 Data Labeling and Annotation Protocol

Annotation of the dataset was conducted using LabelImg software (Tzutalin, 2015), following strict protocols to ensure consistency, accuracy, and high-quality labeling suitable for object detection algorithms. Bounding boxes delineated healthy and diseased parts of banana plants, including leaves, stems, and fruits, following established practices for plant disease detection (Ramcharan et al., 2017).

Annotation protocols required that bounding boxes: (i) be placed when 90% or more of the subject area is visible and not occluded; (ii) encompass all visible edges of the visible subject while maintaining a small margin to provide context around the subject; (iii) capture clearly identifiable mono-infections (Ren et al., 2016).

Expert plant pathologists reviewed annotations and provided feedback on the accuracy of the class labels, resulting in a high-quality dataset preparation and accurate disease identification (Selvaraj et al., 2019). To maintain compatibility with deep learning frameworks for object detection, annotations were saved in PASCAL VOC XML format (Everingham et al., 2010).

#### 2.2.4 CNN Architecture and Model Development

We trained a series of SSDLite MobileNetV2 architectures for banana disease detection. This architecture was selected for its demonstrated speed and success in agricultural applications and plant disease detection, particularly for offline operation on a smartphone (Ramcharan et al., 2017).

SSDLite MobileNetV2 is designed for lightweight, real-time detection, making it ideal for mobile applications and field deployment where computational resources are limited (Howard et al., 2017; Liu et al., 2016).

All models were trained using Azure Machine Learning Services (MLS) on 1 NVIDIA Tesla V100 GPU with 112 GB of system RAM and a 6-core processor. The training was done using TensorFlow’s Object Detection API with transfer learning, leveraging pre-trained weights from the Microsoft COCO dataset to improve feature representation and reduce training time (Huang et al., 2017; Lin et al., 2014). Model performance was evaluated during training using mean Average Precision (mAP) with an IoU threshold of ≥0.5, providing standardized metrics for assessing detection accuracy (Padilla et al., 2020).

#### 2.2.5 Continuous Model Improvement Framework: Integration of Farmer Field Data and Machine Learning Operations Workflow (MLOps)

Building on the dataset collection, preprocessing, and model implementation outlined above, a robust MLOps pipeline was incorporated to continuously improve the AI diagnostic system through iterative real-world data validation and expert feedback **(Figure 1)**, ensuring scalable deployment across diverse agricultural environments (Paleyes et al., 2022; Sculley et al., 2015).

**Figure 1.**
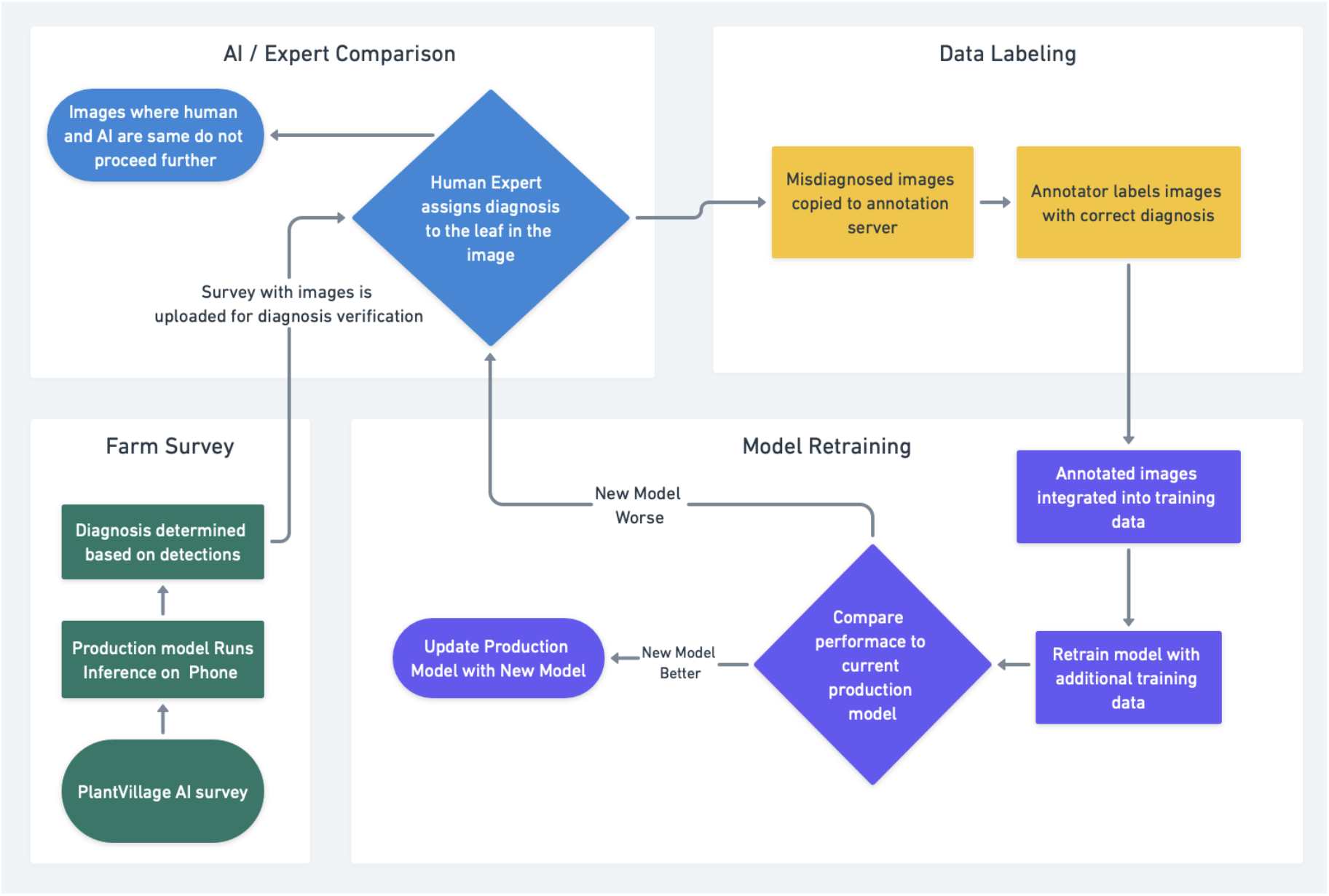
Continuous Model Improvement Framework: Integration of Field Deployment, Expert Validation, and Iterative Retraining for Enhanced Diagnostic Accuracy

##### Farmer Field Deployment and Data Collection

The trained model was integrated as a feature into the PlantVillage Android app to analyze banana plant images and provide real-time diagnoses, along with management advice, directly to farmers in the field (Hughes & Salathe, 2016). Images were streamed directly from the smartphone’s camera to the model for on-device processing, ensuring functionality in areas with limited or no internet connectivity. When connectivity was available, the survey data, including captured images and AI diagnoses, were uploaded to the PlantVillage AI Observatory platform for validation and continuous model improvement.

##### Validation and Error Correction: AI Observatory and Annotation Server

In the PlantVillage AI Observatory, expert plant pathologists systematically reviewed field-collected images to validate the accuracy of AI model predictions (Sahiner et al., 2019). Correctly diagnosed images were confirmed and marked as such, whereas misdiagnosed samples were flagged for expert re-annotation and integration into the next training dataset. These flagged images were forwarded to trained annotators, who applied established protocols to assign correct classifications. The correctly annotated images were integrated into the following training dataset, creating a continuously expanding and improving training dataset for model optimization (Paleyes et al., 2022).

##### Model Retraining and Performance Evaluation

The augmented dataset was used to train new versions of AI models through Azure Machine Learning Services (MLS). Newly trained models were deployed to “staging” versions of the PlantVillage app, accessible only to internal testing teams. If the retrained model outperformed the production model in internal testing, the production model was replaced with the newly trained model. Otherwise, the process returned to expert validation for further refinement and data collection (Kazmierczak et al., 2024).

##### Iterative Improvement and Scalability

This continuous improvement framework enabled the AI diagnostic system to evolve and improve through iterative feedback loops, leveraging real-world field data and expert annotations to enhance accuracy and robustness across diverse environmental conditions **(Figure 1)**. The integration of field-deployable AI with centralized validation and iterative retraining provides a scalable solution for banana disease detection across diverse agroecological zones, while aligning with established best practices in plant disease diagnostics and MLOps implementation (Arpteg et al., 2018; Sculley et al., 2015).

## 3.0 Results

### 3.1 LAMP Assay Development and Performance Evaluation

#### 3.1.1 LAMP Primer Design and Validation

Four LAMP primers (F3, B3, FIP, BIP) were designed to target the DNA-S coat protein gene of BBTV, specifically nucleotides 456–805 bp, using the NEB LAMP Primer Design Tool (**Figure 2**). Primer design was informed by multiple sequence alignment of 33 BBTV isolates from diverse African geographical regions (**Table 1**), ensuring that primers targeted conserved functional domains within the coat protein gene and diagnostic applicability across the genetic diversity of BBTV variants circulating throughout Sub-Saharan Africa

**Figure 2.**
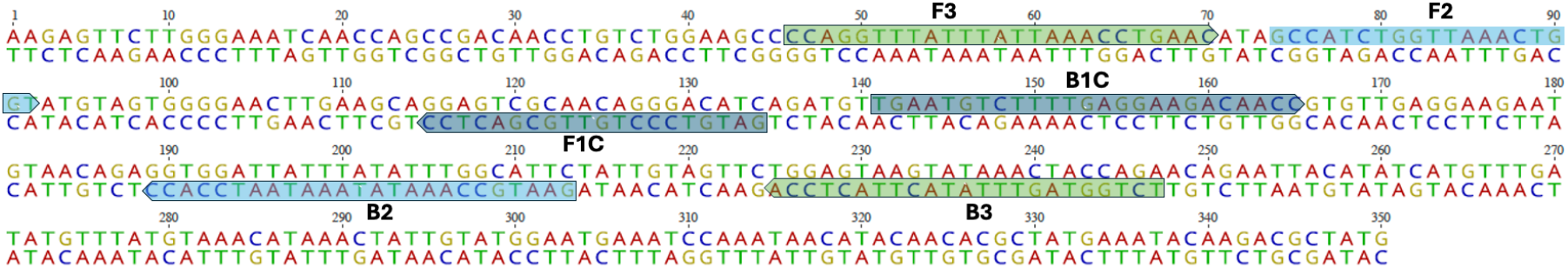
Primers’ positions on the targeted region of DNA-S

#### 3.1.2 Experimental Validation Using Field Samples

Field validation studies were conducted using samples collected from Buhigwe, comprising banana stems and leaves from BBTV-infected plants. The LAMP assay demonstrated specificity, with fluorescence signals consistently exceeding established detection thresholds and being observed exclusively in BBTV-positive samples during the 60-minute isothermal amplification period at 65°C. Notably, both conventional DNA-extracted samples and those prepared using the simplified LAMP extraction buffer protocol yielded comparable fluorescence signals (**Figure 3**). Comprehensive negative-control testing demonstrated the assay’s specificity. Extraction buffer controls, healthy banana leaf samples, and pseudostem tissues consistently showed no detectable amplification signals throughout the entire 60-minute reaction period, with all fluorescence values remaining substantially below the established detection threshold. The assay achieved 100% specificity, with no detectable cross-reactivity with host plant DNA.

**Figure 3.**
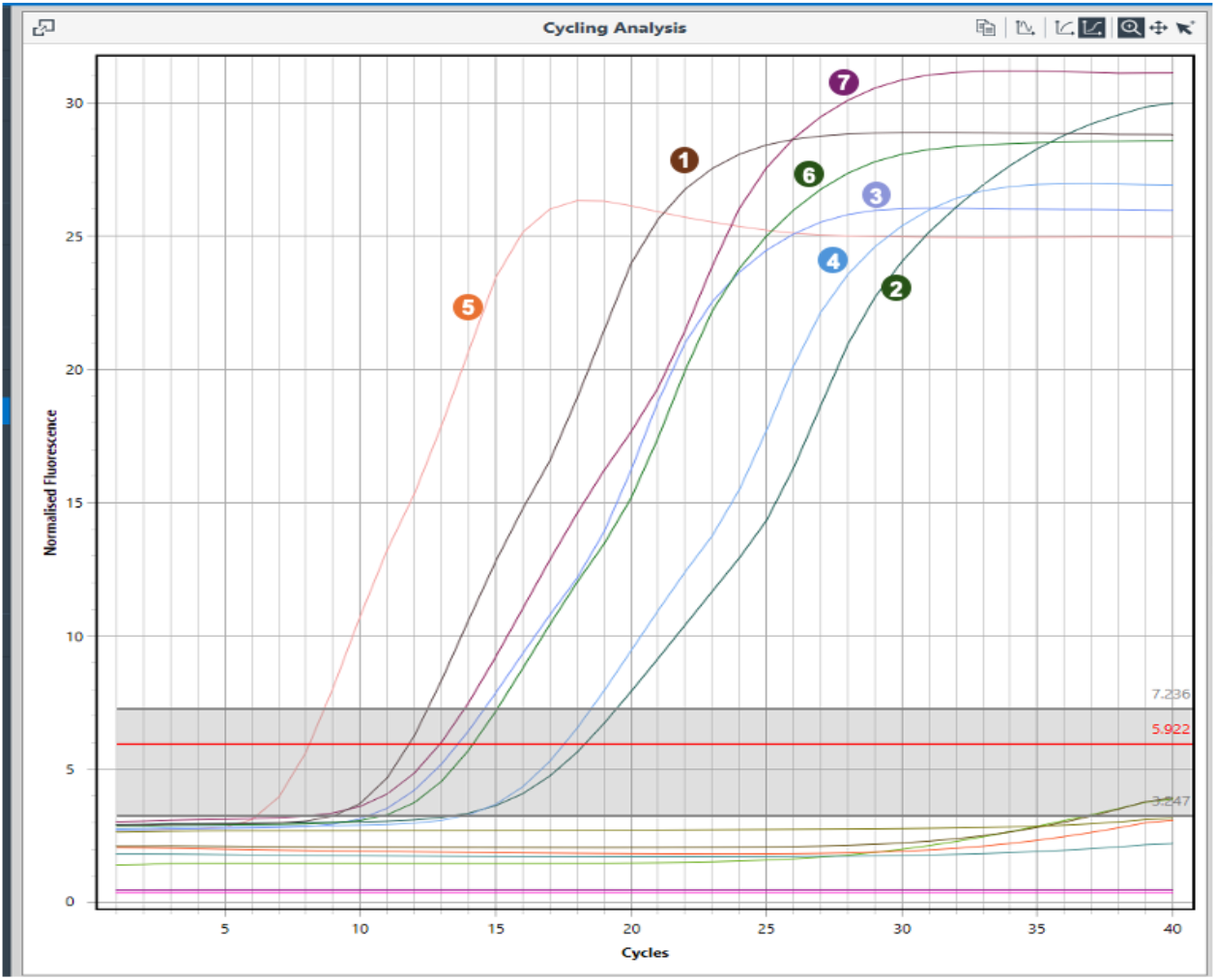
Primers evaluation of LAMP test Fluorescence: (1) DNA extracted from the leaf of an infected plant; (2, 3, 4) single leaf discs from three individual infected plants collected from farmers’ fields and processed with the extraction buffer; (5, 6, 7) pseudostem pieces from three individual infected plants collected from farmers’ fields and processed with the extraction buffer. No fluorescence above the cut-off value was observed for the negative controls, which included DNA from a healthy plant, extraction buffer alone, and leaf and pseudostem samples from healthy plants processed with the extraction buffer.

#### 3.1.3 Comparative Performance of Purified Bst LF Polymerase

The performance characteristics of purified recombinant *Bst* LF polymerase were evaluated through systematic comparative analysis with commercial NEB *Bst* 2.0 DNA Polymerase in parallel LAMP reaction configurations. Using identical BBTV-positive sample sets and comprehensive negative control panels, the comparative analysis revealed that, although the commercial enzyme initiated exponential amplification at approximately 35 minutes, the purified recombinant enzyme showed a consistent 6-minute delay, with amplification onset at approximately 41 minutes (**Figure 4A**). Despite this temporal offset in reaction kinetics, both enzymes achieved comparable endpoint fluorescence intensities of approximately 85-90 fluorescence units and exhibited parallel exponential growth phases, confirming equivalent amplification efficiency and maintaining identical specificity profiles. Specificity evaluation of the purified *Bst-*LF polymerase demonstrated discrimination between BBTV-infected DNA and non-target sequences: positive control replicates exhibited strong fluorescence signals, whereas negative controls using healthy banana DNA and water showed no detectable amplification, confirming the absence of cross-reactivity and false-positive amplification (**Figure 4B**).

**Figure 4.**
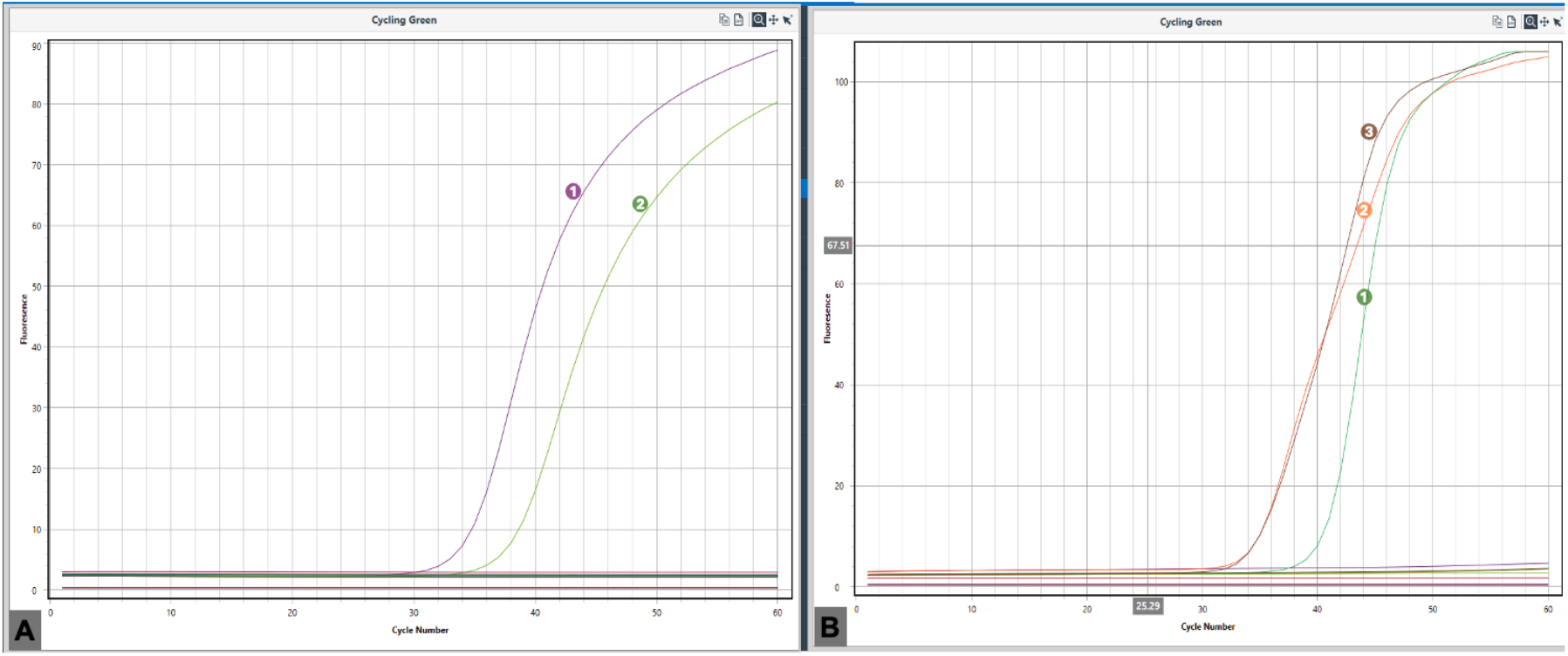
Performance and Specificity Assessment of Purified BstLF Polymerase. **(A)** Comparison of amplification efficiency between purified *Bst*-LF polymerase (Curve 1) and commercial *Bst*-LF polymerase (Curve 2) using BBTV-infected plant DNA as the positive control and healthy banana DNA as the negative control. The fluorescence intensity curves demonstrate successful amplification of BBTV DNA by both polymerases, confirming the activity of the purified enzyme. **(B)** Specificity evaluation of purified *Bst*-LF polymerase in distinguishing BBTV-infected DNA from non-target DNA, including healthy banana DNA and water. Positive control curves (1, 2, and 3) represent replicates using BBTV-infected DNA, showing fluorescence. Negative control curves for healthy banana DNA (4, 5, and 6) and water (7, 8, and 9) exhibit no fluorescence, confirming the absence of cross-reactivity and false-positive amplification.

#### 3.1.4 Comparative Analysis with Conventional Molecular Methods

A systematic comparative evaluation of the LAMP assay against conventional PCR and qPCR was conducted using identical sample sets (**Figure 5**). All three molecular diagnostic platforms yielded consistent amplification results: BBTV-positive samples showed detectable signals, whereas negative controls showed no amplification across all platforms.

**Figure 5.**
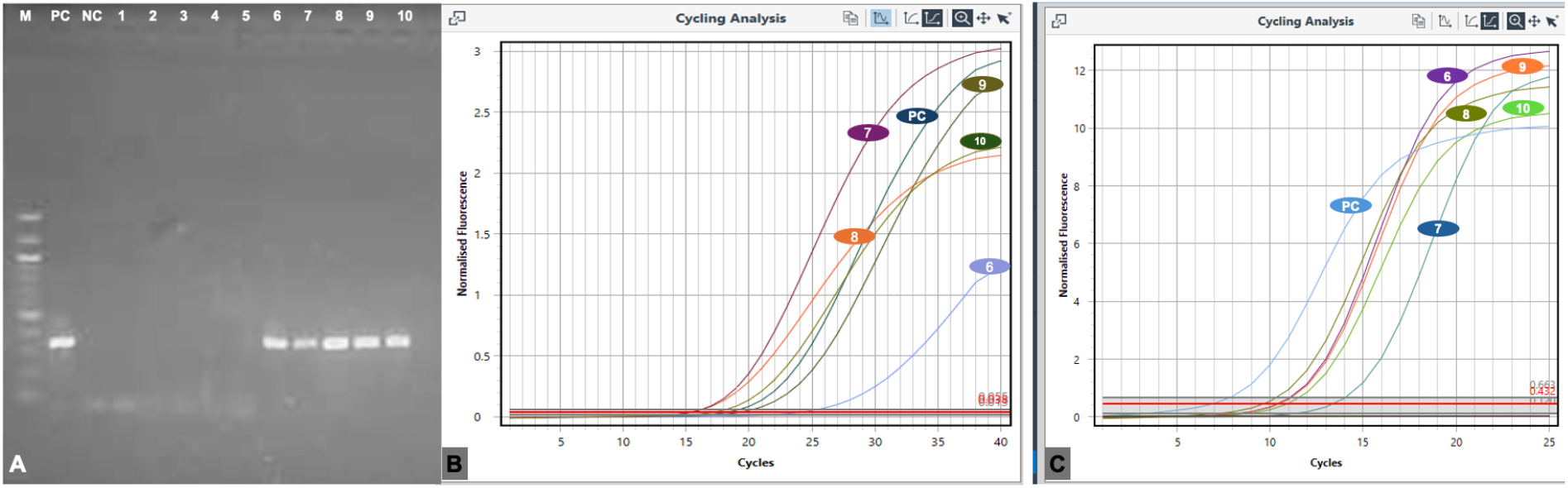
Comparative Evaluation of the Newly Developed LAMP Assay with PCR and qPCR Using Identical Sample Sets. **(A)** Gel electrophoresis results from conventional PCR using samples 6–10 (BBTV-positive) and negative controls (negative control-NC, healthy banana DNA, and water). Clear amplification bands are observed for BBTV-positive samples (positive control-PC and 7–10), while negative controls exhibit no bands, confirming the assay’s specificity. **(B)** Amplification curves from qPCR using the same sample set (6–10, BBTV-positive). Fluorescence signals are detected for BBTV-positive samples (PC and 6–10), while negative controls (NC, healthy banana DNA, and water) show no amplification, confirming the reliability of qPCR for BBTV detection. **(C)** Amplification results from the LAMP assay using the identical sample set (6–10, BBTV-positive). Real-time fluorescence signals are observed for BBTV-positive samples (PC and 6–10), while negative controls (healthy banana DNA and water) exhibit no amplification. The LAMP assay demonstrates comparable performance to PCR and qPCR, while offering operational advantages such as simplified sample preparation (without DNA extraction), real-time visualization, and minimal equipment requirements.

Performance metrics analysis revealed significant operational advantages favoring the LAMP assay: sample preparation time was dramatically reduced from 2-3 hours (conventional PCR and qPCR methods) to under 5 minutes, total assay completion time decreased from 4-6 hours to 60 minutes, and equipment requirements were simplified to single-temperature incubation at 65°C with portable fluorescence detection capabilities.

### 3.2 Computer Vision-Based Disease Detection System

#### 3.2.1 Dataset Development and Model Architecture Evolution

A total of 19 iterations of the optimization process, incorporating systematic dataset refinement and architectural enhancement strategies, were conducted to develop the current diagnostic model (Table 2). The final dataset contained 22 diseases and physiological stress classes, including BBTV, BXW, Black Sigatoka, Yellow Sigatoka, Tattered Leaf, Nutrient Deficiency, and dry leaf conditions, as well as healthy tissue, and was distributed across multiple plant anatomical structures (leaves, pseudostems, buds, and fruits). The initial model iterations (versions 1.0–3.0) used foundational datasets comprising BXW and healthy images of leaves, cut fruits, and cut pseudostems to assess the feasibility of training an AI for banana disease detection. Version 11.0 included additional foliar classes for Black Sigatoka and Yellow Sigatoka.

**Table 2.** Key Developmental Phases in Banana Disease Model Evolution.

| **Development Phase** | **Version** | **Dataset Size** | **Architecture** | **Key Improvements** | **Diseases** |
| --- | --- | --- | --- | --- | --- |
| **Initial Development & Proof of Concept** | 1.0–3.0 | 2,153–2,253 | SSDLite<br>MobileNetV2 | Proved BXW symptoms could be recognized and distinguished from healthy leaves, fruits, and stems | BXW and Healthy |
| **Sigatoka Expansion & Optimization** | 4.0–11.0 | 951–4,998 | SSDLite<br>MobileNetV2 | Added Yellow Sigatoka, Black Sigatoka foliar classes; improved BXW detection with diverse data | BXW, Healthy, Black Sigatoka and Yellow Sigatoka |
| **BBTV Integration & Refinement** | 12.0–18.0 | 9,862–17,736 | SSDLite<br>MobileNetV2 | Major expansion with BBTV whole-plant images; targeted false positive/negative reduction | BXW, Healthy, Black and Yellow Sigatoka, BBTV |
| **Nutrition deficiency Addition & Complete Framework** | 19.0 | 19,914 | SSDLite<br>MobileNetV2 | Nutrient deficiency integration; young plants (sucker) images added | BXW, Healthy, Black and Yellow Sigatoka, BBTV, Nutrient Deficiency |

The BBTV leaf class was introduced in version 12.0, and from then until version 18.0, the dataset was systematically expanded through the methodology described in Section 2.2.5. Finally, the nutrient deficiency class was introduced in version 19. The final optimization in Version 19.0 achieved the most comprehensive disease-detection framework, with 19,914 images across 22 classes, including seven distinct diseases, physiological stress conditions, and healthy tissue categories.

#### 3.2.2 Model Performance Evaluation and Error Analysis

Confusion matrix analysis of the final model (Version 19.0) provided a comprehensive evaluation of diagnostic performance across all 22 classes, revealing both robust classification capabilities and systematic misclassification patterns that informed ongoing dataset refinement strategies (**Figure 6**).

**Figure 6.**
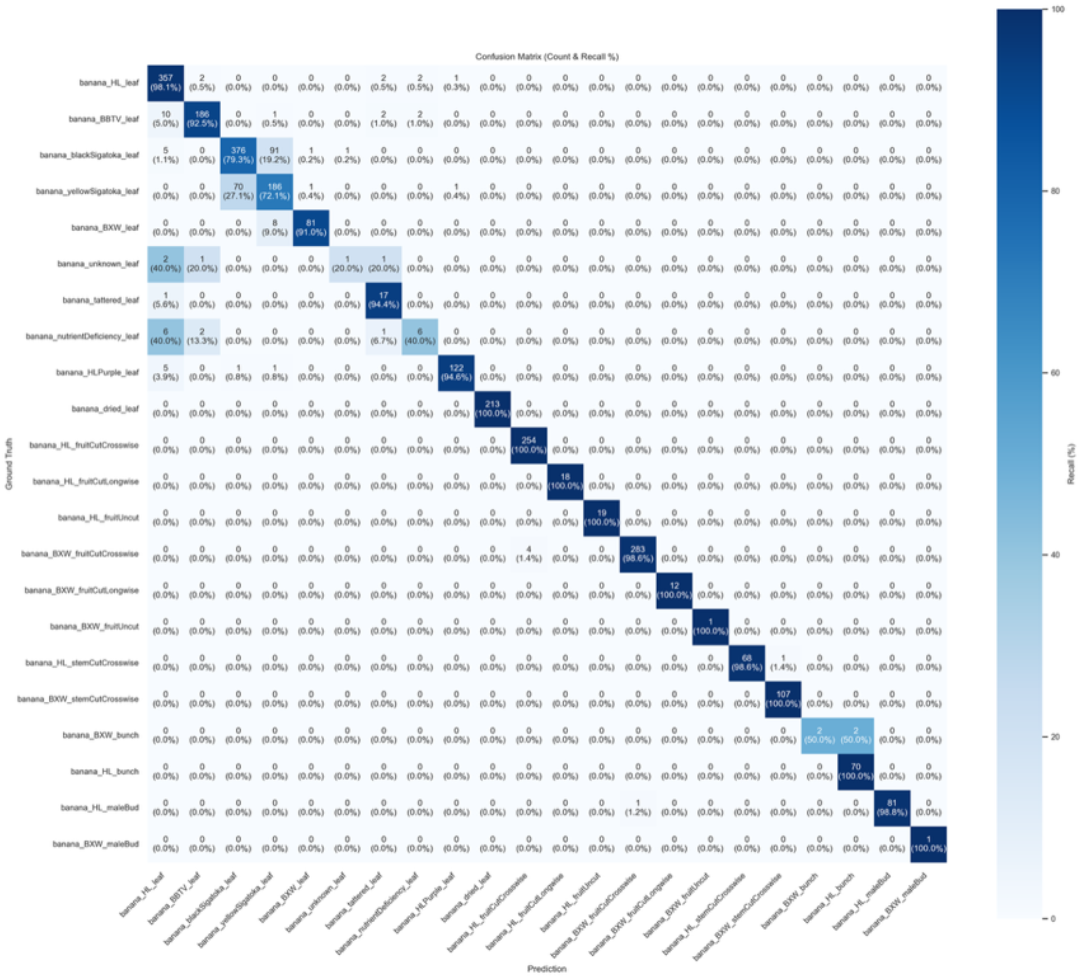
Confusion Matrix Analysis of the Final AI Model (Version 19.0): Classification Performance Across 22 Banana Disease, Physiological Stress, and Healthy Tissue Classes.

The model demonstrated high diagnostic accuracy across most classes. Among foliar disease categories, BBTV leaf detection achieved a recall of 92.5% (186/201 correctly classified), with the primary misclassification pathway involving 5.0% of BBTV samples predicted as healthy leaves. BXW leaf detection attained 91.0% (81/89), with the predominant error being 9.0% misclassified as Yellow Sigatoka. Healthy leaf classification achieved 98.1% (357/364), confirming the model’s strong baseline discrimination capability. The Healthy Purple leaf class, representing young plants exhibiting varietal-derived purple coloration, achieved 94.6% (122/129), indicating that the targeted addition of 710 young-plant images in Version 18.0 effectively resolved earlier false-positive issues in which normal purpling was misidentified as disease symptoms.

The most notable classification challenge occurred between Black Sigatoka and Yellow Sigatoka, reflecting the phenotypic overlap inherent to these closely related foliar diseases. Black Sigatoka achieved 79.3% (376/475), with 19.2% of samples misclassified as Yellow Sigatoka. Conversely, Yellow Sigatoka achieved 72.1% (186/258), with 27.1% misclassified as Black Sigatoka. This bidirectional confusion between the two Sigatoka diseases is consistent with the biological reality that advanced stages of both diseases converge toward similar necrotic leaf phenotypes, and was partially addressed during earlier iterations through the removal of highly advanced symptomatic images that were no longer representative of the characteristic traits distinguishing the two diseases (CHURCHILL, 2010; George et al., 2022).

The model exhibited near-perfect classification performance for dried leaf achieved 100% recall (213/213). Tattered leaf achieved 94.4% recall (17/18). All healthy fruit categories (fruit cut crosswise, fruit cut longwise, and fruit uncut) achieved 100% (254/254, 18/18, and 19/19, respectively). BXW fruit cut crosswise achieved 98.6% (283/287), and BXW fruit cut longwise and BXW fruit uncut both achieved 100% (12/12 and 1/1, respectively). Stem classification was similarly robust, with healthy stem cut crosswise achieving 98.6% recall (68/69) and BXW stem cut crosswise achieving 100% recall (107/107). Healthy bunch and healthy male bud classes achieved 100% and 98.8% (70/70 and 81/82), respectively. BXW male bud achieved 100% (1/1). Two classes exhibited lower performance attributable to limited sample representation in the evaluation set. Nutrient deficiency leaf achieved 40.0% (6/15), with 40.0% of samples misclassified as healthy leaf and 13.3% as Black Sigatoka, suggesting that the visual symptoms of nutrient deficiency overlap substantially with both healthy tissue appearance and early Sigatoka lesions. BXW bunch achieved 50.0% (2/4), though the small evaluation sample size (n = 4) limits the interpretability of this metric.

The iterative error-correction approach implemented between Versions 12.0 and 18.0 was important in achieving the final performance profile. Field testing during this period identified 408 BBTV false negatives and 230 Black Sigatoka false negatives, which were correctly annotated and integrated into subsequent training datasets. Additionally, BBTV false-positive cases were corrected by replacing erroneous labels with correct classifications. The introduction of new classes (Tattered Leaf and Nutrient Deficiency) in Version 18.0 addressed previously unclassified physiological stress conditions that had contributed to false positive Sigatoka and BBTV detections in earlier model iterations. This systematic error-correction process, which relies on expert plant pathologists’ review of misclassified images to generate high-quality ground-truth annotations, demonstrates the critical importance of integrating domain expertise in developing robust agricultural diagnostic systems.

#### 3.2.3 Mobile Implementation and Real-Time Performance

The PlantVillage Android app was updated with a “Banana AI” diagnostic feature that uses TensorFlow 1.12 to deploy a trained Convolutional Neural Network object-detection model. The mobile implementation used SSDLite MobileNet V2 to enable real-time identification of disease symptoms and physiological stress across multiple banana plant parts, including leaves, stems, inflorescences, and fruit structures, with the diagnostic performance described above at Version 19.0. (**Figure 6**).

#### 3.2.4 Integration Framework Implementation

A QR code-based tracking system was implemented to facilitate seamless integration of phenotypic data from Machine-Learning AI model diagnosis with genotypic data from molecular diagnostic platforms. The comprehensive system embedded extensive metadata, including precise GPS coordinates, banana variety specifications, agroecological zone classifications, soil nutrient composition profiles, and meteorological data for each analyzed sample (**Figure 8**).

**Figure 8.**
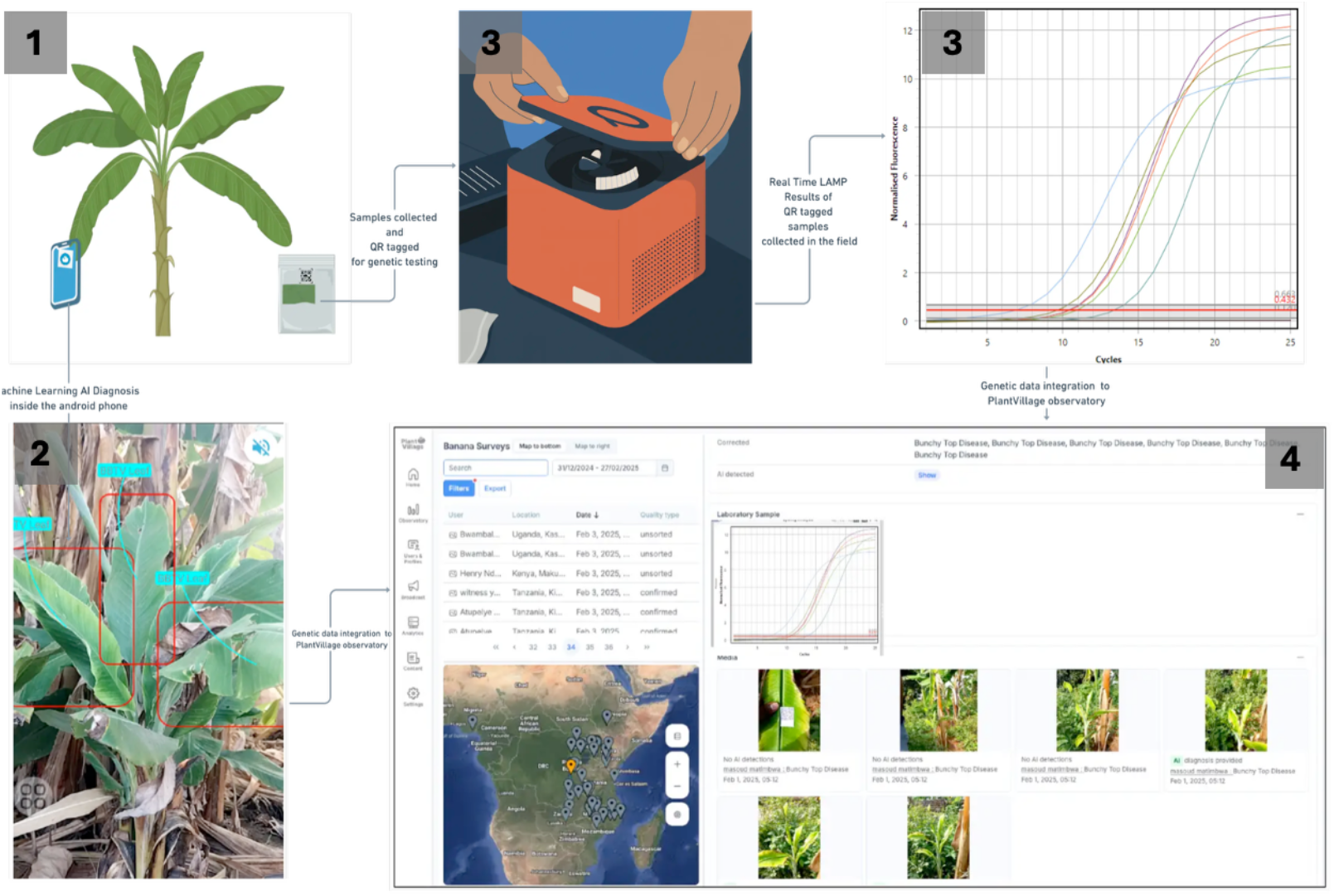
Integrated Diagnostic and Surveillance Framework: Systematic Workflow Combining AI-Powered Phenotypic Screening, LAMP Molecular Confirmation, and Centralized Data Integration for Comprehensive Banana Disease Management **(1)** Field-Based AI Phenotypic Assessment: Smartphone-based computer vision analysis enables real-time disease detection with QR code-based sample tracking for subsequent molecular confirmation. **(2)** Integrated Field Deployment: Extension workers conduct systematic surveys capturing AI diagnoses alongside comprehensive metadata (GPS coordinates, environmental parameters, agronomic practices), with data transmitted to the PlantVillage AI Observatory for expert validation and iterative model refinement. **(3)** LAMP Molecular Diagnostics: Field-collected samples undergo simplified extraction and isothermal amplification, providing definitive molecular confirmation of viral infections, particularly asymptomatic BBTV in planting material. **(4)** Centralized Data Integration Platform: The observatory consolidates phenotypic diagnoses, molecular results, and geospatial data into a unified surveillance architecture, enabling interactive disease distribution mapping and longitudinal epidemiological analysis that transforms isolated diagnostic events into actionable regional disease management intelligence.

The integration framework enabled direct correlation analysis between visual symptoms identified by machine-learning AI assessment and molecular-level pathogen confirmation from LAMP assay results.

## 4.0 Discussion

### 4.1 LAMP Assay Innovation and Diagnostic Advancement

The successful development of LAMP primers specifically targeting the BBTV coat protein gene represents an advance in addressing limitations of existing diagnostic approaches for BBTV in sub-Saharan Africa. The four-primer system’s reduced operational complexity compared with conventional six-primer systems, while maintaining exceptional specificity, aligns with foundational principles (Németh & Kovács, 2025; Notomi et al., 2000) and demonstrates that strategically simplified designs can achieve reliable viral detection in complex field sample matrices. The strategic selection of conserved sequences across 33 BBTV isolates from diverse African regions ensures broad geographic applicability, hence addressing the genetic diversity challenges that have historically limited previous diagnostic approaches (Bouwmeester et al., 2023; Kumar et al., 2011).

We demonstrated that both conventional DNA-extraction samples and crude sample preparations yield comparable fluorescence signals, representing a transformative advancement for field deployment accessibility by eliminating complex DNA-extraction protocols and addressing fundamental time and equipment barriers that have historically prevented the widespread adoption of molecular diagnostics in developing regions (Panno et al., 2020). The ability to obtain reliable diagnostic results within 60 minutes with simplified sample preparation positions LAMP technology as a cornerstone for on-site disease detection.

The assay’s demonstrated specificity profile, with no cross-reactivity with host plant DNA, effectively addresses a pervasive challenge in plant pathogen diagnostics that frequently leads to false-positive results and compromised diagnostic confidence. This specificity performance parallels other successful LAMP assays developed for plant virus detection, which typically demonstrate sensitivities comparable to or exceeding conventional PCR methodologies while offering significant operational advantages (Augustine et al., 2020; Panno et al., 2020).

### 4.2 Economic Sustainability and Scalability Framework

The successful production and validation of purified recombinant *Bst* LF polymerase for the present diagnostic system represents an important breakthrough in addressing economic barriers to large-scale diagnostic deployment. Given that commercial DNA polymerases typically account for 30-40% of total LAMP reaction costs, in-house enzyme production is a critical factor for implementing sustainable diagnostic programs in developing regions, particularly during disease outbreaks (de Souza et al., 2023).

The economic implications extend substantially beyond immediate cost reduction to enable frequent monitoring and testing activities required for effective viral disease outbreak management and to enhance proactive disease management strategies, particularly for outbreaks such as BBTV, where early detection capabilities are fundamental for preventing widespread transmission through infected planting material.

Previous studies show more than 90% savings in polymerase costs when using an in-house *Bst*-LF (Macauyag, 2024), and implementation within this system is projected to lower overall per-reaction costs by 70–80%. These reductions materially improve the feasibility of systematic surveillance and outbreak containment in smallholder farming systems by enabling broader and more sustained testing.

### 4.3 Computer Vision Model Development and Performance

The iterative development process, through 19 systematic iterations, demonstrates the critical importance of continuous optimization for creating diagnostic tools that function reliably in complex agricultural environments. The model detected 22 distinct classes across disease categories, healthy tissue classifications, and physiological stress conditions, representing one of the most extensive diagnostic frameworks developed for banana production constraints in Sub-Saharan Africa (Selvaraj et al., 2019).

The final model achieved a 92.5% true positive rate (TPR) for BBTV detection, with minimal false-positive and false-negative rates. This performance level is particularly significant given BBTV’s devastating impact on banana production and the critical role of early identification in effective disease management.

#### Expert-Informed Error Correction Framework

The systematic resolution of misclassification cases, particularly between BBTV symptoms, healthy young tissue, and nutrient deficiency manifestations, through iterative expert review and dataset refinement highlight the essential role of domain expertise in developing robust agricultural diagnostic systems (Paleyes et al., 2022; Sculley et al., 2015). This expert-informed approach ensured that model improvements reflected actual biological distinctions rather than spurious correlations in training data.

### 4.4 Mobile Implementation and Diagnostic Democratization

#### Technological Optimization for Field Deployment

The mobile implementation framework, which utilizes TensorFlow’s optimization architecture, addresses a fundamental barrier to diagnostic technology adoption in resource-constrained agricultural settings: the need for offline functionality in regions with limited or unreliable internet connectivity (Sandler et al., 2018).

#### Platform Accessibility and Distribution Framework

The diagnostic system has been operationalized through the PlantVillage mobile application, which is publicly available on the Google Play Store (PlantVillage, 2018), thereby establishing a scalable distribution mechanism for the widespread dissemination of technology. This implementation strategy leverages existing mobile infrastructure and user familiarity with application-based platforms, hence reducing adoption barriers associated with specialized hardware requirements or technical training prerequisites.

#### Deployment Viability Across Heterogeneous Technological Environments

The demonstrated performance characteristics, integrating rapid inference processing, offline operational capacity, and cross-device compatibility, position this diagnostic system as uniquely viable for large-scale deployment across technologically heterogeneous agricultural environments (Mohanty et al., 2016b; Ramcharan et al., 2019). This multidimensional optimization addresses an empirically documented phenomenon in which technical complexity and infrastructure requirements constitute primary impediments to adoption, even when diagnostic tools demonstrate theoretical availability and proven efficacy (Hughes & Salathe, 2016; Ramcharan et al., 2019).

#### Implications for Technology Democratization

The combination of diagnostic accuracy, computational efficiency, and broad device compatibility represents a critical advancement toward democratizing access to precision diagnostic technologies within smallholder agricultural systems. By eliminating dependencies on specialized equipment, continuous internet connectivity, and high-performance computing infrastructure, this implementation framework fundamentally transforms the accessibility for AI-powered disease diagnostics in developing agricultural contexts.

### 4.5 Integration Framework and Surveillance System Innovation

The embedded metadata system incorporates comprehensive environmental and agronomic data, addressing the biological reality that disease expression and progression patterns are influenced by complex interactions among pathogen genetics, host genetics, and prevailing environmental conditions (Arpteg et al., 2018).

The ability to integrate these multifactorial influences through data collection provides unprecedented insights into disease ecology and the optimization of management. This integrated approach is particularly valuable for BBTV management, in which the virus can remain latent for extended periods before symptom expression. Under these circumstances, molecular confirmation is essential for accurate assessment of disease status, as phenotypic screening alone cannot reliably detect asymptomatic infections in planting material (Čarija et al., 2022; Ramcharan et al., 2019). The generation of high-resolution disease distribution maps enables evidence-based resource allocation and the implementation of targeted intervention strategies. This capability effectively transforms disease management paradigms from reactive approaches (responding to visible outbreaks) to predictive approaches that anticipate and prevent disease spread. These integrated surveillance capabilities represent a significant advancement over current surveillance systems that lack the data architecture necessary for comprehensive epidemiological analysis (Bouwmeester et al., 2023).

### 4.6 Implications for Sustainable Banana Production Systems

The synergistic integration of molecular diagnostics for the detection of asymptomatic infections with Machine-Learning AI-powered phenotypic screening effectively addresses critical gaps in current banana disease management strategies. Evidence from vegetatively propagated crop systems demonstrates that molecular detection enabling deployment of virus-free planting material can prevent yield losses ranging from 40-90%, depending on crop system and disease pressure (Chikoti et al., 2019; Legg et al., 2017; Namanda et al., 2019).

This prevention capacity is particularly significant given the rapid spread potential of viral diseases via infected planting material and vector transmission pathways (Blomme et al., 2017; Kumar et al., 2011). The economic value proposition of integrated surveillance has been empirically quantified in Australian banana production systems, where modeling studies predicted that BBTV exclusion and early detection strategies prevent annual losses of AUD $15.9-27.0 million, yielding a benefit-cost ratio of 1.8:1.0, demonstrating that every dollar invested in surveillance returns $1.80 in prevented production losses (Cook et al., 2012). These economic returns underscore the sustainability of comprehensive surveillance investments.

Achieving economic accessibility through cost reduction transforms systematic surveillance from a resource-intensive undertaking accessible only to large-scale operations into a viable strategy for smallholder farming communities. This democratization addresses a fundamental barrier to the implementation of effective disease management at the scales required for meaningful agricultural impact. Furthermore, real-time disease mapping capabilities enable precision-based interventions, including the targeted distribution of virus-free planting material and the implementation of evidence-based quarantine measures (Augustine et al., 2020; Hughes & Salathe, 2016).

### 4.7 Future Directions and Broader Applications

The integrated diagnostic framework established through this research, combining LAMP-based molecular detection with AI-powered phenotypic assessment, provides a methodological foundation for expanding capabilities across multiple pathogen targets and crop production systems. For molecular diagnostics, the validated LAMP primer design methodology and simplified sample preparation protocols are directly transferable to additional viral, bacterial, and fungal pathogens affecting banana and other vegetatively propagated crops, including Banana Xanthomonas Wilt, Fusarium wilt TR4, and emerging viral threats. Concurrently, the computer vision framework supports systematic expansion by incorporating additional disease classes and crop species into training datasets, thereby establishing a comprehensive diagnostic ecosystem that transcends single-pathogen limitations while maintaining the cost-effectiveness essential for resource-limited settings.

Critical research priorities include systematic validation across diverse agroecological zones with varying environmental conditions, the development of multiplex LAMP assays that enable simultaneous detection of multiple pathogens from a single sample, and the refinement of field-deployable detection platforms that eliminate dependence on laboratory-based equipment. For Machine-Learning AI-based systems, essential priorities include comprehensive performance validation across additional banana-growing regions with distinct varietal compositions, temporal validation to ensure model robustness across seasonal variations, and the development of active learning frameworks that continuously improve diagnostic accuracy through the systematic integration of field-validated data. Integration protocols must be established for embedding these diagnostic capabilities within existing agricultural extension systems, ensuring that technological innovations translate into accessible decision-support tools for farming communities.

The demonstrated feasibility of deploying sophisticated molecular assays, including in-house polymerase production to enhance economic sustainability alongside smartphone-based AI diagnostics in resource-constrained settings, fundamentally challenges prior assumptions about technological accessibility barriers in developing agricultural regions. This integrated approach is directly extensible to other vegetatively propagated staple crops (cassava, sweet potato, yam) and perennial tree crops (citrus, mango, cocoa), where disease detection in planting material critically influences production outcomes.

